# Autoregulation of cluster root and nodule development by white lupin CCR1 receptor-like kinase

**DOI:** 10.1101/2024.07.04.602037

**Authors:** Laurence Marquès, Fanchon Divol, Alexandra Boultif, Fanny Garcia, Alexandre Soriano, Cléa Maurines-Carboneill, Virginia Fernandez, Inge Verstraeten, Hélène Pidon, Esther Izquierdo, Bárbara Hufnagel, Benjamin Péret

## Abstract

Root development is controlled by local and systemic regulatory mechanisms that optimize mineral nutrient uptake and carbon allocation. The Autoregulation of Nodulation (AoN) pathway defines a negative regulation of nodule development in Legumes as a way to regulate the costly production of nitrogen-fixing organs. This pathway is defined as a response to symbiotic interaction and has been shown to also control root formation to some extent. However, it remains unclear if root and nodule development are under coordinated genetic regulation. Here, we identified mutants with altered root development in white lupin, constitutively producing specialized lateral roots called cluster roots. We showed that the CCR1 receptor-kinase negatively regulates cluster root and nodule development and targets common molecular modules such as NIN/LBD16-NFYA, defining a novel pathway that we named Autoregulation of Development (AoDev). AoDev defines a negative systemic pathway controlling several types of root organ development, independently of symbiotic partners and nutrient availability.

## INTRODUCTION

Plasticity of plant root development enables the exploration and exploitation of soil resources to meet the demand for nutrients. As plant development is essentially post-embryonic, systemic integration pathways play a central role in coordinating plant growth and nutrient acquisition at the whole-organism level. Specifically, the demand for nitrogen (N) triggers a cascade of signaling events that orchestrate systemic responses to optimize nutrient uptake and allocation^1,2^. Recent studies have highlighted the involvement of root-secreted small peptides and their cognate leucine-rich repeat receptor-like kinases (LRR-RLK) as pivotal regulators of local cell-to-cell communication and systemic root-to-shoot-to-root signaling pathways^3,4^. These pathways are of critical importance in both governing plant-wide developmental processes and maintaining nutrient homeostasis, ultimately contributing to optimal plant fitness.

In legume plants, Autoregulation of Nodulation (AoN) has long been recognized as a pivotal negative systemic regulator of nodulation induced by rhizobial colonization^5–7^. Its characterization has been achieved through a combination of EMS-based forward genetics, natural variation approaches and grafting experiments in model species such as *Lotus japonicus*, *Glycine max,* and *Medicago truncatula*^8–11^. Under conditions of nitrate deficiency, legume roots are predisposed to colonization by N-fixing rhizobacteria, resulting in the formation of symbiotic nodules. The AoN pathway encompasses two distinct mechanisms that finely tune the regulation of root nodulation to balance shoot and root development, thereby preventing excessive proliferation of energy-consuming nodules^7,12^. The first mechanism involves the production of CLAVATA3/EMBRYOSURROUNDING REGION-RELATED (CLE) peptides in the roots, followed by their translocation via the xylem to the shoot. Upon reaching the shoot, CLE peptides interact with LjHAR1/GmNARK/MtSUNN LRR-RLK receptors, leading to the inhibition of nodulation. The second mechanism, elucidated in *M. truncatula,* and also responsive to nitrate supply, involves the action of root secreted C-TERMINALLY ENCODED PEPTIDE (CEP) peptides and the MtCRA2 LRR-RLK receptor in shoot tissues. This mechanism not only governs lateral root development but also, unlike the CLE-dependent route, activates nodulation^13^. Recent investigations in *M. truncatula,* have identified the microRNA miR2111, and TOO MUCH LOVE (TML) Kelch-repeat F-box proteins, as components of the downward signaling cascade in both routes^14,15^.

White lupin (WL, *Lupinus albus*) is a hardy legume crop capable of thriving in nitrogen- and phosphate-deficient soils, owing to the remarkable developmental plasticity of its root system^16,17^. In response to N deficiency, white lupin roots establish a symbiotic relationship with *Bradyrhizobium lupini,* resulting in the formation of nitrogen-fixing nodules. Additionally, phosphate starvation triggers the development of specialized lateral roots known as “cluster roots” (CRs)^18^. These CRs are characterized by densely clustered, short third-order lateral roots termed rootlets, which serve as highly efficient organs for nutrient acquisition^19,20^. Through the secretion of protons, organic acids, and phosphatases, CRs facilitate the solubilization of phosphate pools that are otherwise inaccessible to conventional root systems. In contrast, narrow-leaved lupin (NLL, *L. angustifolius*) is another resilient crop species that does not produce CRs^21^. However, they exhibit the capacity to mobilize phosphate pools through enhanced phosphatase activity^22^. Despite the fact that white lupin has been employed as a model for root exsudative activities, it remains unclear how cluster roots are induced and how their development is controlled at the whole plant level.

Here, we employed a forward genetic approach based on screening of a mutagenized white lupin population for enhanced CR production in P-rich conditions, and identified the *Lupinus albus CONSTITUTIVE CLUSTER ROOT 1* (*LalbCCR1*) gene. This gene encodes an LRR-RLK, exhibiting synteny with the LRR-RLKs involved in AoN, namely *LjHAR1, GmNARK and MtSUNN.* CCR1 functions in a systemic pathway that restricts CR numbers, reminiscent of AoN, even in the absence of rhizobial symbiosis and in the presence of nitrogen. We demonstrate that by controlling both CR and nodule numbers, CCR1 is involved in a global root developmental pathway, which prevents the overproduction of root organs that excessively deplete carbon resources. We named this regulatory mechanism Autoregulation of Development (AoDev).

## RESULTS

### Genetic identification of *LalbCCR1* as a LRR-RLK inhibiting CR development

In phosphate-deficient medium (-P), white lupin exhibits numerous CRs in the upper part of the root system, whereas P-rich medium (+P) inhibits their development (Fig. 1a). We screened 800 M2 batches of an ethyl-methanesulfonate (EMS)-mutagenized WL AMIGA population for plants exhibiting numerous CRs in the P-rich suppressive condition. We identified four independent lines of recessive mutants (ems5, ems96, ems120, ems413). These mutants consistently develop two to five times more CRs compared to wild-type plants independently of the Pi supply (Fig. 1a-b; Extended data Fig. 1a). Additionally, they have shorter lateral and primary roots resulting in a marginally lower dry weight compared to wild-type (Fig. 1b; Extended data Fig. 1b). CRs of the mutants display a higher rootlet density, as if all the potential sites of third LR initiation have been unlocked (Fig. 1c-d). Physiologically, the CRs of the mutants are fully active, excreting large amounts of protons, and displaying elevated phosphatase and reductase activities (Extended data Fig. 1c). The pronounced overproduction of CRs observed in the mutants supports the involvement of the mutated gene in an inhibitory pathway of CR development in wild-type plants.

**Figure 1.**
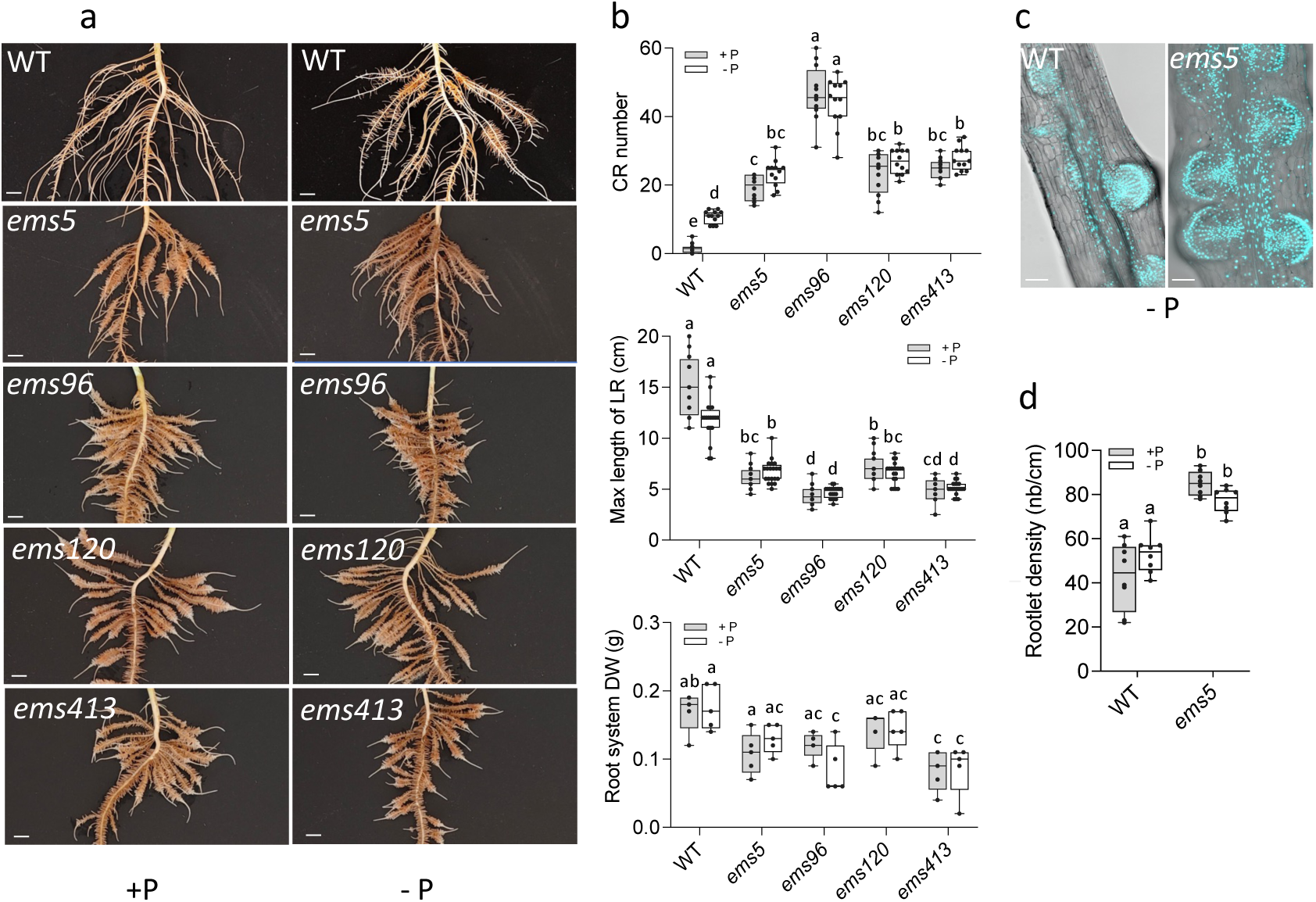
Phenotypes of the four allelic *ccr1* mutants. a) Representative images of the upper part of the root system from 20-day-old wild-type (WT) and the four *ccr1* mutant plants grown on either phosphate-rich medium (+P) or phosphate-deficient medium (-P). Scale bar = 1 cm. b) Quantitative analysis of various root traits in wild-type (WT) and *ccr1* mutant plants, including CR abundance within the upper 10 cm of the root systems, maximum lateral root length, root system dry weight. Statistical analysis was performed using two-way ANOVA with Tuckey correction, p < 0.05. c) Apotome imaging of a CR section from the *ems5* mutant compared to wild-type (WT) plants. DAPI staining revealed rootlet primordia, identifiable by their small nuclei. Scale bar = 100 µm. d) Density distribution of rootlet along 1 cm of cluster root in the *ems5* mutant. Statistical analysis was performed using two-way ANOVA with Tuckey correction, n = 8, p < 0.05

The mutations are recessive, segregate with the mutant phenotype, and the four independent mutants fall into the same complementation class that we named *ccr1* (*constitutive cluster root 1*) (Extended data Fig. 2a-b). Bulk-Segregant Analysis combined with mapping-by-sequencing (BSA-seq) was performed in parallel on the four allelic *ccr1* mutant lines. This analysis unveiled a linkage of causal mutations to the beginning of chromosome 3 in all four *ccr1* lines (Extended data Fig. 2c). Further analysis of this genomic region across the four *ccr1* alleles pinpointed the presence of causal SNPs within the *CONSTITUTIVE CLUSTER ROOT 1* gene (*LalbCCR1, Lalb_Chr03g0025491,* https://www.whitelupin.fr). *LalbCCR1* encodes a putative protein like kinase of the LRR-RLK family XI-1 (Fig. 2a) with a canonical 3D-structure calculated by homology modeling using ColabFold v1.5.5: AlphaFold2 with MMseqs2 (Fig. 2b). Specifically, the *ccr1-2* allele carries a G1370A SNP resulting in a premature stop-codon within the LRR domain; both *ccr1-1* and *ccr1-3* alleles contain a G2099A SNP causing a G700E substitution in the kinase domain; and *ccr1-4* exhibits a C2437T SNP resulting in an H813Y change within the kinase domain (Fig. 2a). Notably, the G>E substitution found in *ccr1-1* and *ccr1-3* has also been identified in a hypernodulating pea line (P91, *Pssym29-7*)^23,24^. This mutation is also present in a *LalbCCR1* paralog, that we named *LalbCCR1-like* (Fig. 2c). The pangenome analysis of WL confirms the presence of this SNP in *LalbCCR1-like* across all WL accessions^25^. Therefore, it is reasonable to infer that LalbCCR1-like is non-functional in WL, resulting in the absence of functional redundancy between LalbCCR1 and LalbCCR1-like, thereby explaining the efficiency of EMS mutagenesis in isolating mutants with a highly constitutive CR phenotype. In order to describe the relationship between *LalbCCR1*, *LalbCCR1-like* and other ortholog genes, a phylogenetic tree was constructed, including ten of the nearest BLAST-sequence homologs of *LalbCCR1* in *Arabidopsis thaliana*, and LRR-RLKs from model legumes known to be involved in AoN: LjHAR1, GmNARK, MtSUNN, and MtCRA2 (Fig. 2d). LalbCCR1 clustered with GmNARK, LjHAR1, MtSUNN, as the nearest LRR-RLK/XI homolog of AtCVL1. Further analysis of genomic loci revealed that the three legume LRR-RLK genes *MtSUNN, LjHAR1, GmNARK* are syntenic to *LalbCCR1* (Fig. 2e).

**Figure 2.**
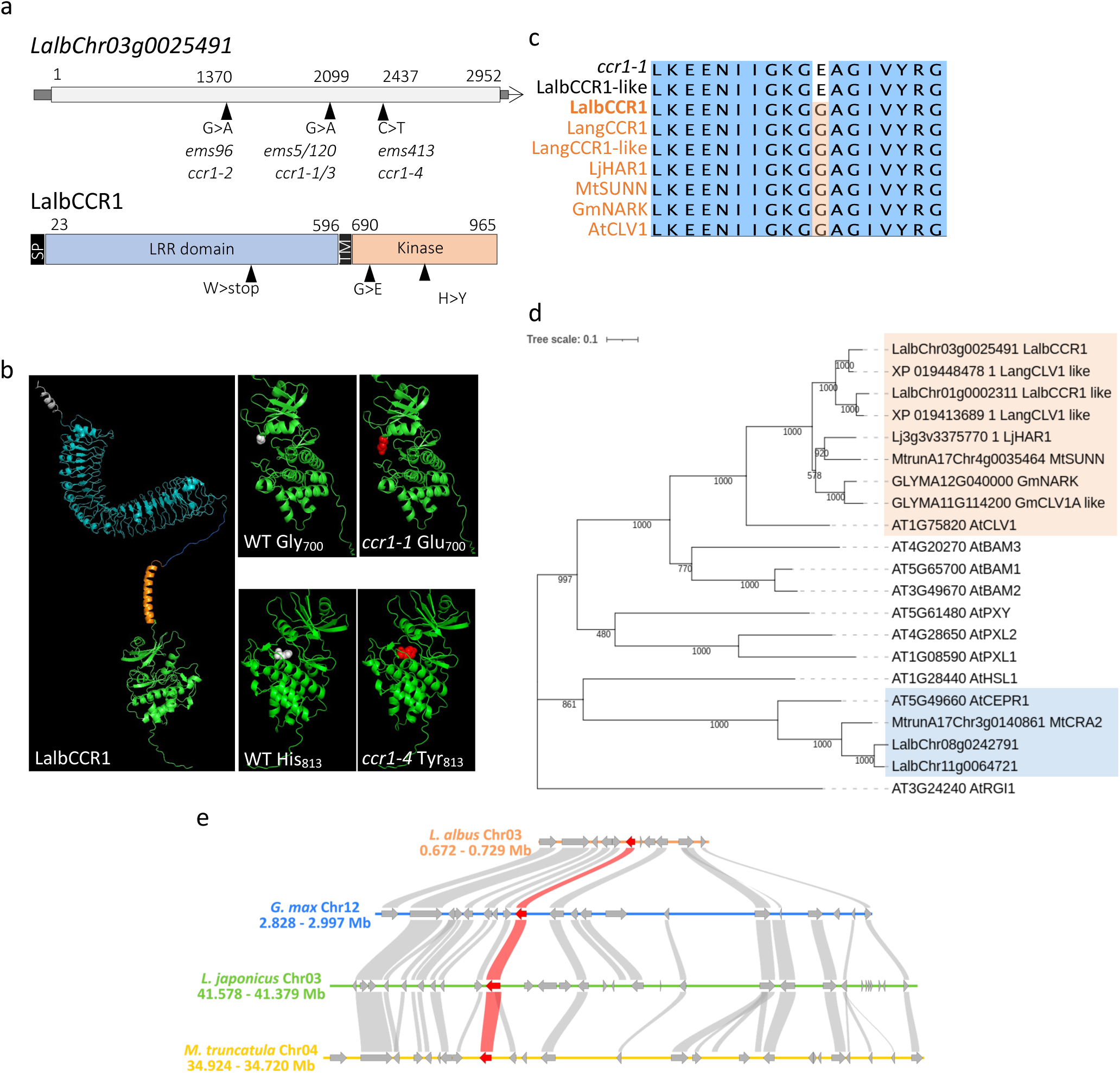
LalbCCR1 is a LRR-RLK syntenic with LRR-RLKs involved in Autoregulation of Nodulation. a) *LalbCCR1* gene (*LalbChr03g0025491*) and LalbCCR1 protein structures. The *LalbCCR1* gene lacks introns. EMS-induced mutations are indicated by a triangle with the corresponding mutant names. The predicted LalbCCR1 protein contains a LRR domain, a transmembrane domain and a kinase domain. The specific amino-acid substitutions resulting from EMS-induced SNPs are given. b) The 3D-structural model of LalbCCR1 generated by AlphaFold2, highlighting the mutated amino-acid positions within the kinase domain of *ccr1-1* and *ccr1-4* mutants. c) Phylogenetic tree of LalbCCR1-related proteins across narrow-leaved lupin (*Lupinus angustifolius*), *Lotus japonicus*, *Glycine max*, *Medicago truncatula* and *Arabidopsis thaliana*. d) Allelic variations in the kinase domain segment containing the *ccr1-1/3* G>A mutation across closest-homologs of LalbCCR1 in *L. albus*, *L. angustifolius, Lotus japonicus*, *Glycine max*, *Medicago truncatula* and *Arabidopsis thaliana.* Notably, the *LalbCCR1* paralog, *LalbCCR1-like* exhibits the same punctual mutation observed in *ccr1-1/3* mutants, leading to a non-functional protein in *L. albus*. e) Syntenic relationships among *LalbCCR1, GlmNARK, LjHAR1, MtSUNN* loci, indicating genomic organization across species at this locus.

### LalbCCR1 controls CR and nodule development through a systemic shoot-to-root signaling pathway

*MtSUNN, LjHAR1* and *GmNARK* are well known to be involved in AoN with corresponding mutants displaying remarkable supernodulation phenotypes^5,11,26,27^. Based on the synteny observed between *LalbCCR1* and these three LRR-RLKs, we assessed the nodulation phenotype of the independent *ccr1-1* and *ccr1-2* mutants after inoculation with *Bradyrhizobium lupini.* Both mutant lines exhibited hypernodulating phenotypes (Extended data Fig. 3a-f). The efficiency of nitrogen fixation was highlighted by the healthier appearance and greener foliage of nodulated plants compared to the non-nodulated ones, as well as by the reddish coloration of the nodules. These hypernodulation phenotypes in *ccr1* mutants provide additional validation for the accurate identification of the causal gene in a species where stable transformation and subsequent complementation assays are not feasible. We then used grafting experiments to test the systemic nature of hypernodulation phenotypes observed in *ccr1* mutants. Our results demonstrated that *ccr1* shoots promote hypernodulation, characterized by clusters of nodules intertwined with short rootlets, when grafted onto wild-type roots (Fig. 3a-c; Extended data Fig. 3g-j). Therefore, *LalbCCR1* governs nodule development from shoots through a systemic shoot-to-root pathway. Analysis of *LalbCCR1* expression in various WL organs revealed predominant expression in petioles and hypocotyls irrespective of P conditions (Extended data Fig. 4). Thus, *LalbCCR1* appears to be the ortholog of *MtSUNN, LjHAR1, GmNARK* for AoN.

**Figure 3.**
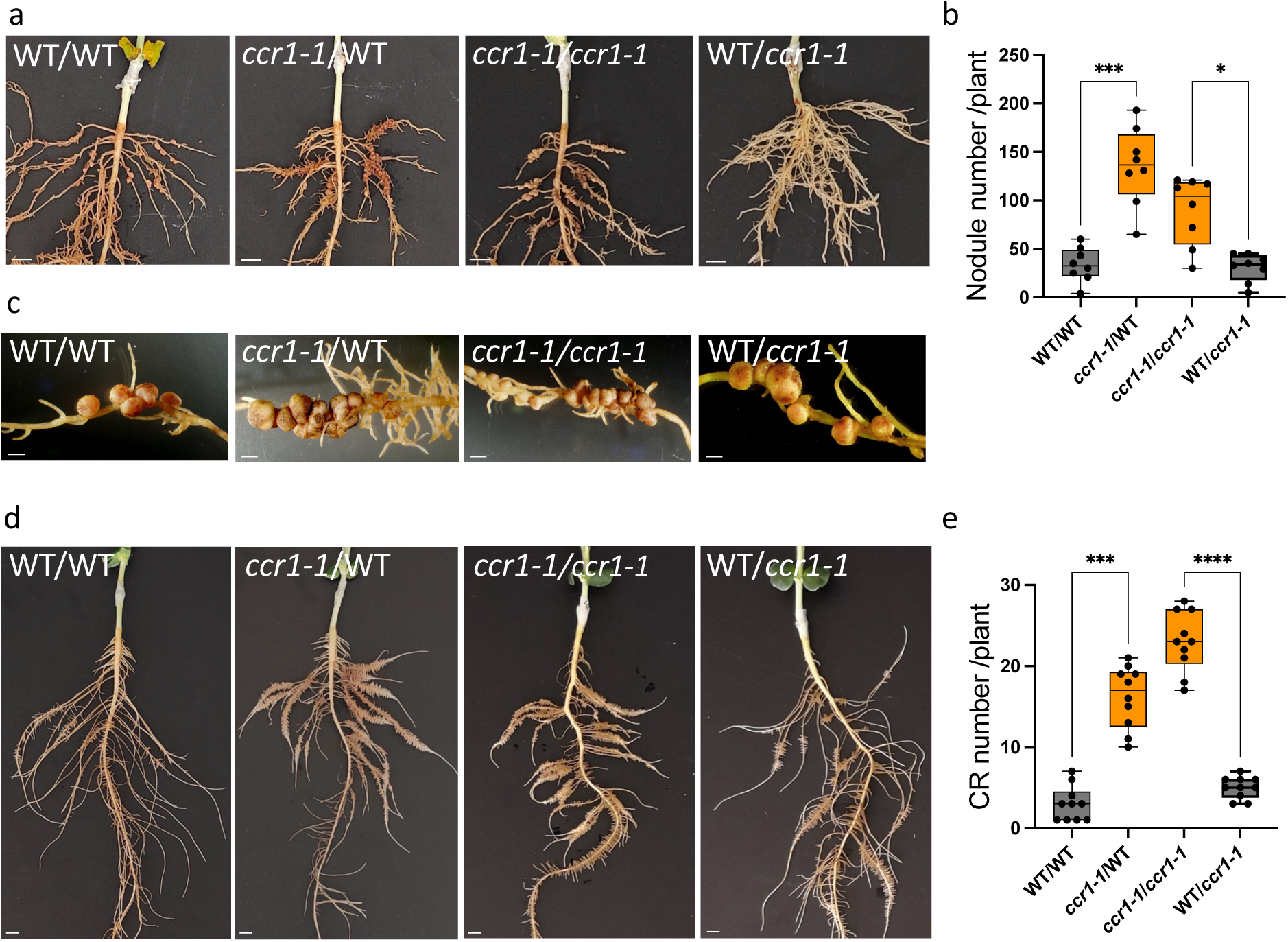
*LalbCCR1* controls CR nodulation and development via a systemic signaling pathway. The four graft combinations involving wild-type (WT) and *Lalbccr1-1* mutant plants were cultivated on either P-rich or N-deprivated medium and inoculation with *Bradyrhizobia lupini* was performed in Magenta boxes, or in tanks containing N- and P-rich conditions for phenotypic evaluation of the root systems. a) Representative images of nodulated rootstocks of the four 3-week-old graft combinations inoculated with *Bradyrhizobia lupini*. The legends of the images indicate scion/rootstock. Scale bar = 0.7 cm. b) Quantification of nodule numbers per plant on the rootstocks of the four 3-week-old graft combinations. The legends of the images indicate scion/rootstock. Error bars represent mean ± SD, with statistical analysis performed using Kruskal-Wallis test, n = 8, *adjusted p-value = 0.0305, ***adjusted p-value = 0.0003. c) Magnification of the nodules formed on the rootstocks of grafted plants, revealing compact clusters of nodules on *ccr1-1*/WT and *ccr1-1*/ *ccr1-1* plants. The legends of the images indicate scion/rootstock. Scale bar = 0.2 cm. d) Root systems of the four 3-week-old graft combinations. The legends of the images indicate scion/rootstock. Heterografted plants, *ccr1-1*/WT and WT/*ccr1-1*, demonstrate that the constitutive CR phenotype is primarily influenced by the *ccr1-1* mutation in the scion rather than in the rootstock. Scale bar = 1 cm. e) Quantitative assessment of CR abundance within the upper 10 cm of the rootstocks. Error bars represent mean ± SD, and statistical analysis was performed using Kruskal-Wallis test, n = 10, ***adjusted p-value = 0.0006, ****adjusted p-value<0.0001.

Subsequently, we conducted grafting experiments to elucidate whether *LalbCCR1* controls CR development via a systemic mechanism akin to AoN. All grafting experiments were conducted under P-rich conditions, which are favorable for the grafting process and known to inhibit CR development. In this CR-suppressing condition, significant CR development occurred when *ccr1* shoots were grafted onto wild-type rootstocks, while conversely, grafting wild-type shoots significantly inhibited CR formation on *ccr1* rootstocks (Fig. 3 d,e and, for *ccr1-2,* Extended data Fig. 5a,b). These grafting outcomes unequivocally demonstrate that *LalbCCR1* governs CR development via a systemic shoot-to-root pathway, reminiscent of AoN. They highlight that *LalbCCR1* drives a common systemic shoot-to-root pathway that regulates both nodule and CR development in WL.

### Identification of a common negative systemic signal across *Lupinus* species

We took advantage of the inability of NLL to form cluster roots to test whether the CCR1-dependent inhibitory signal is conserved across *Lupinus* species. Since WL and NLL possess differing chromosomal numbers precluding interspecific crosses, we performed interspecific grafting experiments. Grafting *ccr1* mutant shoots onto NLL rootstocks induced a remarkable transformation in NLL root architecture, leading to the development of clusters of short tertiary roots, that could be described as CR-like (Fig. 4 a,b and, for *ccr1-2,* Extended data Fig.5 c). This striking modification occurred in P-rich medium, beneficial for grafting, indicating that the P-starvation signal is not involved in this phenotype. Grafting of *ccr1* shoots was thus sufficient to initiate the formation of CR-like structures in a non-CR producing lupin species, suggesting the existence of a potent inhibitory systemic pathway in NLL. We also observed that grafting wild-type WL shoots onto NLL rootstocks, triggered the development of some tertiary short roots, albeit to a lesser extent than when *Lalbccr1* was used as scion. A dose-effect relationship appeared in the induction of CR development of NLL rootstocks across the three grafting scenarios: NLL/NLL, WL/NLL and *Lalbccr1*/NLL, with an increasing proportion of short tertiary roots respectively (Fig. 4a,b). An inverse developmental gradient was observed when *Lalbccr1* was used as rootstock. Grafting wild-type WL shoots onto *Lalbccr1* rootstocks, inhibited CR development, albeit to a lesser extent than when NLL was used as scion (Fig. 4c,d and, for *ccr1-2,* Extended data Fig. 5b,d). Indeed, NLL shoots profoundly and strikingly inhibited the development of CRs on *ccr1* roots. It should be noted that neither *LangCCR1* nor *LangCCR1-like* carry the *ccr1-1* SNP while *LalbCCR1-like* was found to harbor it (Fig. 2d). The strong inhibitory effect of NLL shoots could be explained by the fact that NLL lupin likely possesses two functional LRR-RLK CCR1s whereas WL has only one. These allelic variations in CCR1 and CCR1-like LRR-RLKs between WL and NLL may explain the observed phenotypic gradient in our grafting experiments and should be explored further. This finding underscores the major role played by these LRR-RLKs in regulating root architecture development in *Lupinus spp* through a conserved inhibitory systemic shoot-to-root pathway.

**Figure 4.**
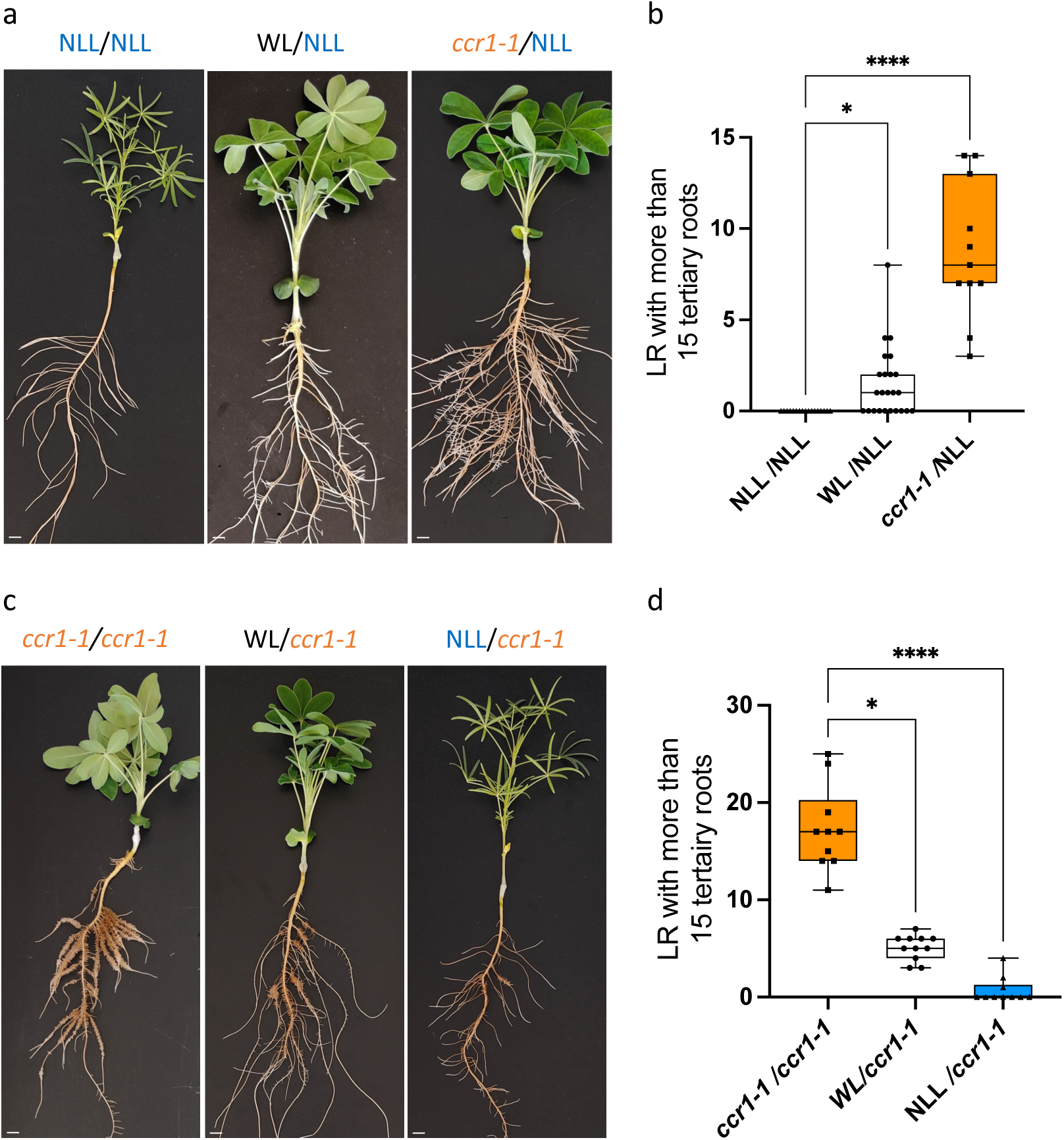
Phenotypes of interspecific grafts between *L. albus*, *L. angustifolius*, and the *ccr1-1* mutant. a) Representative images illustrating rootstocks phenotypes of grafted plants using NLL (*L. angustifolius*) as rootstock and NLL, WL (*L. albus)* or the mutant *Lalbccr1-1* as scions. The *Lalbccr1-1* scion triggered the emergence of numerous tertiary roots on NLL rootstock. Legends of the images indicate the scion/rootstock combination. Scale bar = 1 cm. b) Quantitative evaluation of the abundance of LR with more than 15 tertiary rootlets on NLL rootstocks, with statistical analysis performed using Kruskal-Wallis test; n = 11 to 24; adjusted p-values are indicated: *p = 0.0110, ****p<0.0001. c) Representative images illustrating rootstocks phenotypes of grafted plants using the *Lalbccr1-1* mutant as rootstock and NLL *(L. angustifolius),* WL (*L. albus) or Lalbccr1-1* as scions. The NLL scion markedly suppressed the formation of CRs on *Lalbccr1-1* rootstock, whereas WL did so to a lesser extent. Notably, *LalbCCR1-like* gene carried the same mutation as in the *Lalbccr1-1* mutant, whereas *LangCCR1-like* does not. Legends of the images indicate the scion/rootstock combination. Scale bar = 1 cm. d) Quantitative evaluation of the abundance of LR with more than 15 tertiary rootlets on *Lalbccr1-1* rootstocks, with statistical analysis performed using Kruskal-Wallis test, n = 10 to 11, adjusted p-values are indicated : *p = 0.013, ****p<0.0001.

### Lateral root developmental genes are involved in early CR development

White lupin CR serves as an excellent biological model for investigating tertiary roots development, as it undergoes successive emergence of numerous rootlets along one lateral root (LR), establishing a continuous spatial and temporal gradient of developmental stages. Taking advantage of this model, we conducted sampling of rootlet formation by collecting 1 cm-long root segments every 12h, up to 132h as described before^28^ to generate a comprehensive temporal transcriptomics dataset of cluster root development. Additionally, pieces of LRs were utilized as control (Fig. 5a; Supplementary Table 1). Principal component analysis of sample distribution delineates an elliptical trajectory progressing from state T000, proximal to LR, to timepoints T024-036-048 (hours), which exhibit the greatest deviation from LR samples along axis 1. These three T024-036-048 samples are not distinctly differentiated, unlike the more consistent states observed at T120 and T132, when rootlets had completed their growth (Fig. 5b). In order to identify genes specifically expressed during the early developmental stages of rootlets, we retrieved 2349 differentially regulated genes between LR and timepoints T024-036-048 (Absolute Log2(fold change) > 2; FDR < 0.05) (Supplementary Table 2). Subsequent Gene Ontology (GO) enrichment analysis highlighted terms associated with “cell division”, “cell cycle”, “response to auxin” and “root development”, among the up-regulated genes with the most prevalent counts indicating a clear alignment with our objective to target the early developmental stages of rootlet formation (Fig. 5c; Supplementary Table 3). Conversely, GO enrichment analysis of down-regulated genes resulted in a less interpretable outcome, with numerous GO terms spanning diverse pathways (Extended data Fig. 6; Supplementary Table 3). Following filtration for up-regulated transcription factors, we obtained 144 genes (Supplementary Table 4), and we compiled a list of transcription factor genes up-regulated from 8 families known to be involved in root development: AP2/EREB, LOB domain, ARF, GRAS, Homeodomain, NAM, PLATZ and STY-LRP1 (Supplementary Table 5). We retrieved 55 WL genes, with the AP2/EREB family representing nearly half with a total of 27 genes. We employed the “Orthologous gene search” tool available on our WL website (https://www.whitelupin.fr), combining the Orthologous Matrix Algorithm (OMA) and NCBI BLAST searches to retrieve genes homologous to *Arabidopsis thaliana*. Within the AP2/EREB family, we discovered genes associated with ethylene and cytokinin signaling, such as ERF and CRF, alongside well-characterized genes implicated in early LR patterning, including 4 PUCHI, 2 AIL6/7, and 3 PLT1/2 genes. Additionally, within other families, we identified genes crucial for LR development, including LBD16, LBD29, WOX5, LRP1, SMB, SCR (Fig. 5d). The temporal expression patterns of these genes align closely with well-established transcriptional dynamics observed during *A. thaliana* LR development^29^, providing compelling evidence that rootlets undergo analogous developmental processes as LR (Fig. 5e) and pinpointing sets of early and dynamic gene expression patterns as potential targets of the AoDev pathway.

**Figure 5.**
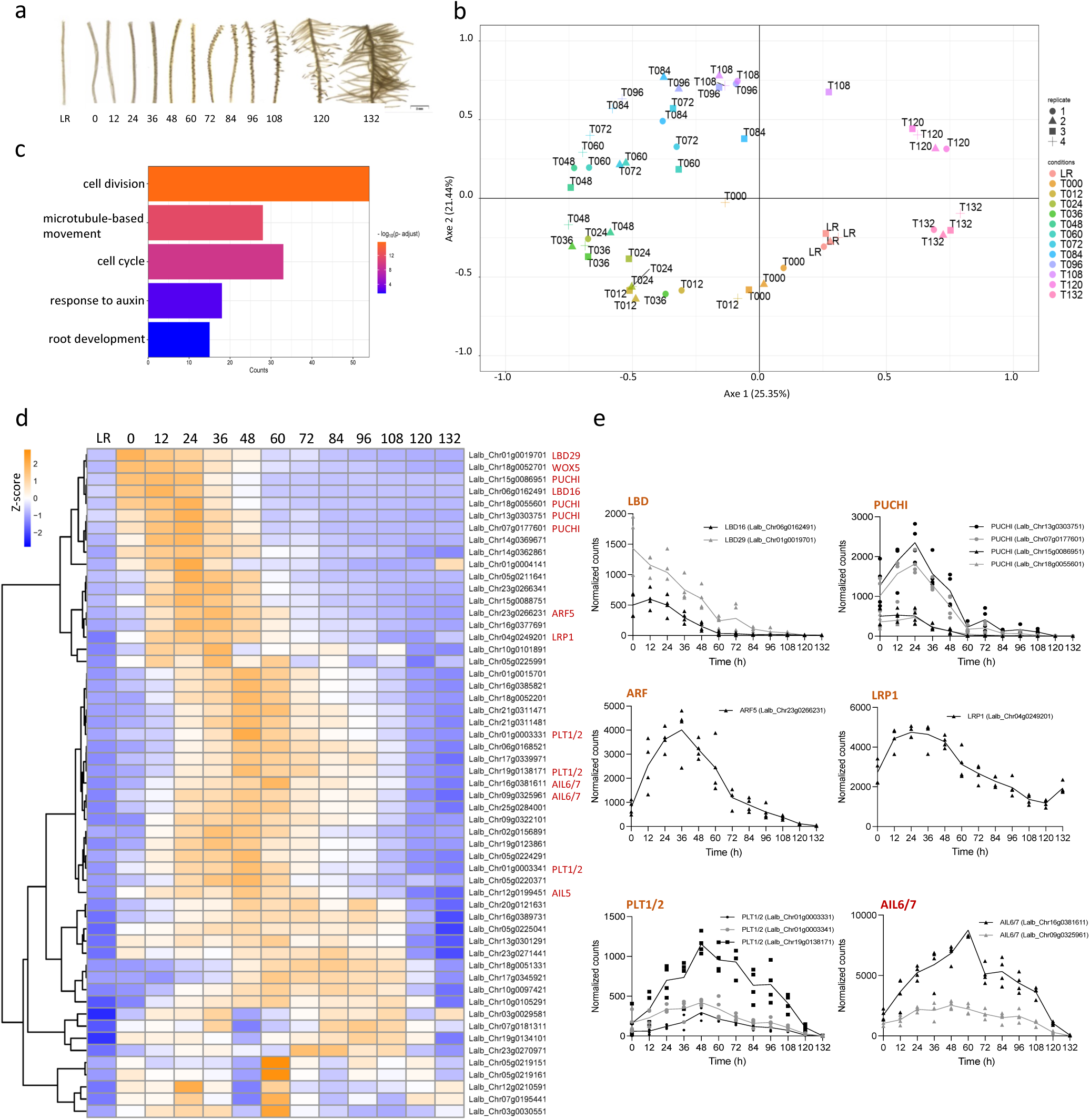
Transcriptional expression analysis during CR formation in white lupin. a) Illustrations depicting the developmental stages of 1-cm long segments of LR sampled for developmental temporal RNAseq transcriptomic analysis. Samples were collected every 12h over a period of 132h. Segments of LR without developing rootlets served as control. Four biological replicates were performed, each containing eight 1-cm-long LR segments coming from four different plants. Scale bar = 2 mm. b) Scatter plot presenting the distribution of the RNAseq dataset with the four replicates for each timepoint and LR control samples in the two principal components, explaining 46.79% of the total variance. c) GO enrichment analysis of early up-regulated genes between LR and timepoints T024-036-048 (Absolute Log2(fold change) > 2, FDR < 0.05). GO terms presenting counts > 10 are displayed. They are associated with cell division, response to hormones and root development. Normalization, differential expression and GO term analysis were performed using DIANE R package^57^. d) Clustered heatmap displaying the normalized Z-scores of gene expression for up-regulated transcription factors from 8 families known to be involved in root development (AP2/EREB, LOB domain, ARF, GRAS, Homeodomain, NAM, PLATZ and STY-LRP1), comparing LR with timepoints T024-036-048 (Absolute Log2(fold change) > 2, adjust p-value (FDR) < 0.05). Transcription factors known to participate in *A. thaliana* LR patterning are highlighted. e) Kinetic profiles illustrating changes over the 132h early developmental stages of six major LR patterning transcription factors involved in CR development.

### Both developmental and *NIN* genes are up-regulated in *ccr1* plants independently of rhizobial infection

To elucidate the regulatory role of *LalbCCR1* in CR development and identify potential targets of AoDev, we performed a RNAseq transcriptomic analysis focusing on LR transitioning into CR, at timepoints close to T024-036-048 of prior RNAseq data, in wild-type and *ccr1-1* mutant plants grown in P-deficient medium (Fig. 6a; Supplementary Table 6). This growing condition was chosen to induce CR formation in wild-type plants and reveal early dynamic changes rather than on/off responses. Differential gene expression analysis between wild-type and *ccr1-1* (Absolute Log2(fold change) > 1; FDR < 0.05), revealed 1845 deregulated genes, with 798 up-regulated and 1047 down-regulated genes (Supplementary Table 7). GO enrichment analysis revealed the activation of oxidative and phosphate stress response pathways in the *ccr1* mutant, highlighting its ability to perceive and respond to phosphate deprivation (Extended data Fig. 7a; Supplementary Table 8). Nevertheless, it is noteworthy that the *PHR1* genes are not up-regulated in *ccr1* plants compared to wild-type, suggesting that the phosphate starvation response is not differentially engaged in the mutant line compared to wild-type plants. Interestingly, a significant up-regulation of several CLE and CEP peptides was observed, indicating a widespread deregulation of the AoN signaling within the mutant line, consistent with previous results. Analysis of down-regulated genes did not retrieve relevant outcomes (Extended data Fig. 7b). Further filtering for up-regulated genes identified 28 transcription factors that were shared with the up-regulated transcription factors in early stages of CR development (Fig. 6b, Supplementary Tables 5 “Common gene in the Venn diagram”). Notably, transcription factors such as LBD16, LBD29, WOX5, PUCHI, AIL6/7, and PLT1/2 were found. These results were confirmed in another *ccr1* allele by measuring gene expression levels on LRs transitioning into CRs from *ccr1-2* plants by RT-qPCR (Extended data Fig. 7c). Altogether, using two complementary transcriptomics approaches, we revealed that genes expressed early during CR development and known to be important regulators in other models for LR development (such as LBD16) are up-regulated in the *ccr1* mutants.

**Figure 6.**
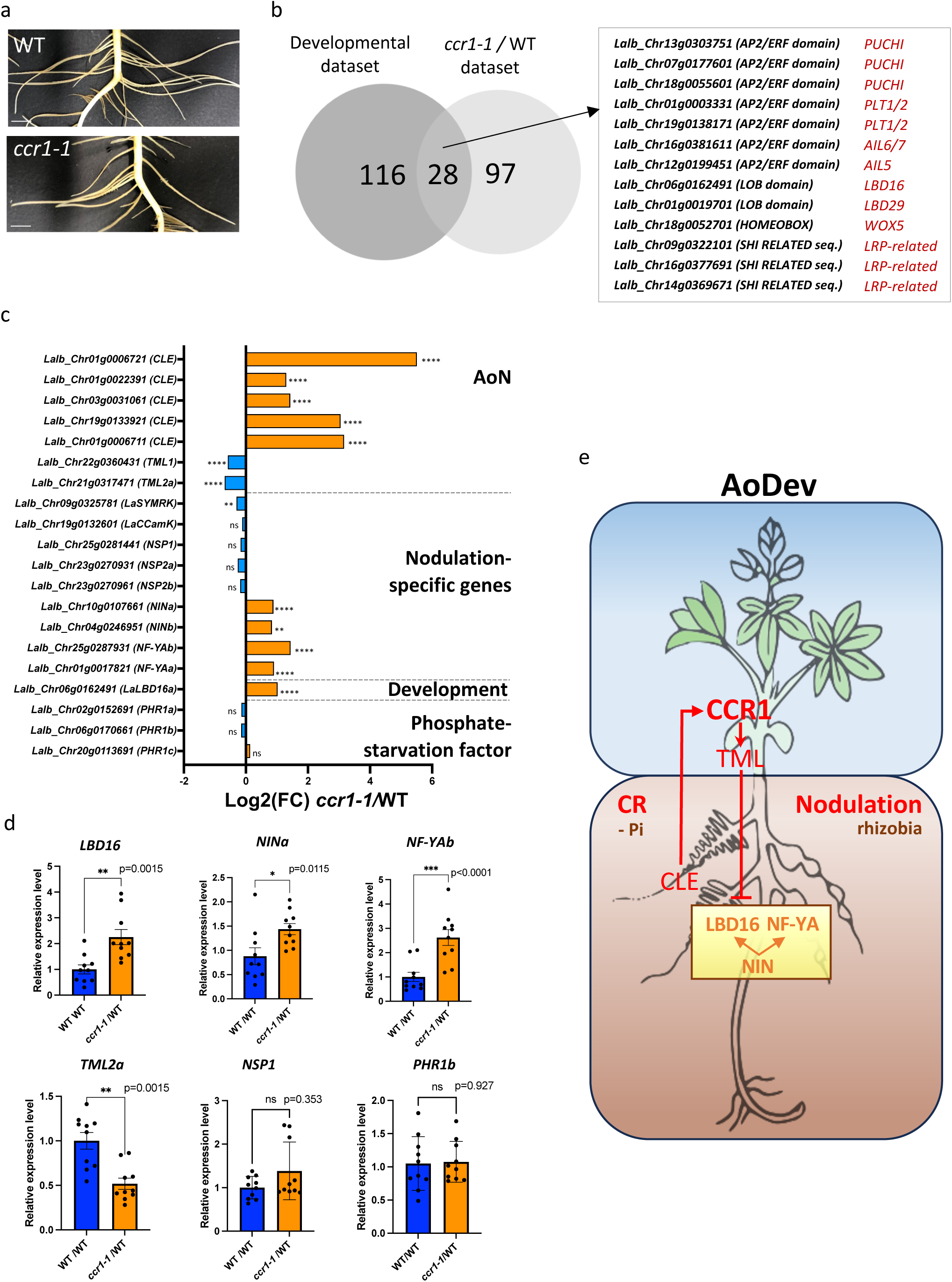
Transcriptional expression analysis of *ccr1-1* mutant during CR development and the proposed model for the AoDev pathway. a) Illustrations depicting the developmental stages of roots used for sampling the LRs from WT and *ccr1-1* plants for the RNAseq. LRs longer than 3 cm from three plants were pooled, and four biological replicates were collected. Scale bar = 1 cm. b) Venn diagram illustrating the overlap of transcription factors-encoding genes up-regulated at timepoints T024-036-048 (hours) versus LR samples in the temporal developmental RNAseq dataset (minimal gene count sum across conditions = 1200, Absolute Log2(fold change) > 2, FDR < 0.05) compared to those up-regulated in the *ccr1-1* CR versus wild-type samples (minimal gene count sum across conditions = 80, Absolute Log2(fold change) > 1, FDR < 0.05). Normalization and differential expression analysis were performed using DIANE R package^57^. Among the 28 shared genes, 13 emblematic transcription factors involved in LR/CR initiation are listed. c) Log2(fold change) of selected key genes in the *ccr1-1* mutants relative to wild-type plants, focusing on the AoN, the nodulation pathway, and the *LBD16* and *PHR1* genes. Expected regulations were observed for AoN genes with up-regulation of *CLE* genes and down-regulation of *TML* genes. Nodulation genes were not significantly deregulated in *ccr1-1* mutant compared to wild-type, except for *NIN* and *NF-YA. PHR1* genes were not significantly deregulated in *ccr1-1* mutant compared to wild-type. Fold changes and adjusted p-values (FDR) were calculated using DIANE R package^57^. In the graph: ns p-adjust>0.05, *p-adjust<0.05, **p-adjust<0.01, ***p-adjust<0.001, ****p-adjust<0.0001. d) RT-qPCR analysis of the expression levels of selected genes in wild-type rootstocks of grafted plants using either wild-type (WT/WT) or *ccr1-1* mutants (WT/*ccr1-1*) as scions. The grafting experiment revealed that *LBD16*, *TML*, *NIN*, *NF-YA* genes are systemically regulated by *LalbCCR1* via a shoot-to-root systemic pathway. Additionally, it confirmed that the *NSP1* nodulation gene or *PHR1* genes are not significantly deregulated when comparing WT and *ccr1-1* in the growing conditions used. Error bars represent mean ± SE, and statistical analysis was performed using Two-tailed Mann Whitney tests: ns p-value>0/05, *p-value <0.05, **p-value <0.0, ***p-value <0.00, ****p-value<0.0001. e) Proposed model for the AoDev pathway, depicting a systemic root-to-shoot-to root signaling mechanism involving the LRR-RLK CCR1, which represses the *NIN/LBD16/NF-YA* module, leading to the inhibition of both CR and nodule development.

Analyzing the deregulated genes between *ccr1-1* mutant and wild-type plants, we observed that *CLE* and *TML* genes exhibited the expected regulatory patterns corresponding to their role in AoN : up-regulation for CLE and down-regulation for TML (Fig. 6c). However, unexpectedly, we found that two *NIN* genes and two *NF-YA* genes, specific to the symbiotic pathway, were slightly but significantly up-regulated in *ccr1-1* plants compared to wild-type, while other genes involved in the nodulation pathway, including *SYMRK*, *CCamK*, or *NSP1*, did not show significant up-regulation in *ccr1-1* background (Fig. 6c; Supplementary Table 9). Additionally, *PHR1* genes did not exhibit up-regulation in *ccr1-1* mutants compared to wild-type. RT-qPCR expression analysis of a second *ccr1* allele (*ccr1-2)* validated these regulations for *NIN*, *NF-YA, NSP1 and PHR1* (Extended Fig. 7d). Given the expression of *LalbCCR1* in both roots and shoots, we investigated whether the observed regulation of *NIN* and *NF-YA* was due to local or systemic events. Grafting experiments were conducted, and LRs transitioning into CRs were sampled from recovering roots. Expression analysis revealed that *LBD16*, *NIN* and *NF-YA* genes were up-regulated in wild-type rootstocks when *ccr1* mutants were used as scions, while *TML*, *NSP1* or *PHR1* were not (Fig. 6d). These findings suggest that in wild-type plants, a shoot-to-root systemic pathway involving *LalbCCR1* leads to the direct or indirect inhibition of *NIN* gene expression, which may control both *LBD16* and *NF-YA* expression independently of nodulation and *PHR1* pathways. Therefore *LalbCCR1* regulates genes involved in both cluster root and nodule development in a shoot-to-root control, defining the Autoregulation of Development pathway.

## DISCUSSION

The root system of White Lupin exhibits a robust developmental plasticity, producing nodules under conditions of nitrogen deficiency in the presence of compatible rhizobia, and CRs in the event of phosphate deprivation. Specifically, CR development has been poorly studied and lacks genetic and molecular description, due to the absence of established model species among the numerous species producing these structures (mainly trees and shrubs, mostly perennials). We took advantage of the recently published genome and pangenome in white lupin^25,30^ to conduct a forward genetic screen and identified four allelic *ccr1* mutants consistently producing CRs, irrespective of phosphate availability. Independent mapping of the 4 mutants pinpointed *LalbCCR1* as the causative gene responsible for inhibiting CR development in wild-type plants. Strikingly, *LalbCCR1* encodes an LRR-RLK sharing synteny with the CLV1-like LRR-RLKs well-established in legume AoN^8,9,31^. We demonstrated that LalbCCR1 governs both AoN and CR development through a systemic shoot-to-root pathway, indicating the involvement of the same systemic pathway in both CR formation and nodule organogenesis. This observation prompted us to define an Autoregulation of Development (AoDev) pathway that encompasses the regulation of both structures (Fig. 6e).

Recent evidence indicates the incorporation of part of the LR developmental program into root nodule development in leguminous plants^32,33^. The Nodule Inception (NIN) transcription factor plays a pivotal role in this integration, serving as a key regulator that bridges the two programs by modulating the expression of both *LBD16* and *NF-YA* genes^34,35^. In our study, transcriptomic analysis of CR development highlighted the critical role of LBD16 as a transcriptional factor essential for CR initiation, akin to its importance in LR patterning^36^. Subsequently, we conducted another transcriptomic analysis comparing young wild-type and *ccr1-1* LR, grown in a phosphate-deficient medium. Our analysis revealed the deregulation of several genes associated with AoN, such as *CLE, CEP* and *TML,* as expected. Meanwhile intriguingly, we also observed up-regulation of *NIN* and *NF-YA* genes in *ccr1* mutants compared to wild-type, despite the absence of rhizobial infection and nodule organogenesis. Canonical genes involved in rhizobia infection, such as *SYMRK*, *CCamK*, or *NSP1/2* did not show any up-regulation, consistent with the absence of rhizobia. On the other hand, *PHR1,* a transcription factor known to be induced upon phosphate deficiency and to up-regulate *NIN* expression^37^, did not exhibit an up-regulation in *ccr1-1* compared to wild-type samples. Therefore, this P-starvation pathway cannot be responsible for the local increase in *NIN* expression in the *ccr1-1* roots. These two observations, namely up-regulation of the *NIN/LBD16-NFYA* module and the absence of regulation by rhizobial and nutrient (phosphate) signals in the *ccr1* mutant, advocate for a developmental level of regulation. The AoDev pathway hereby targets shared molecular components to refrain from organ formation.

Grafting experiment, backed up by qPCR analysis performed on LR rootstocks, definitively established the systemic nature of the up-regulation of *NIN/LBD16-NFYA* module, confirming the differential expression patterns observed in the RNAseq dataset. This systemic regulation is significant, as leguminous plants exhibit intricate local and systemic regulatory networks involving AoN LRR-RLKs to modulate responses to nutrient availability, particularly for N and P supply^38^. For instance, in soybean, rhizobial infection triggers a systemic root-to-shoot-to-root inhibition of nodulation via the RIC1/2-NARK module, while the same NARK receptor can locally suppress nodulation through the production of NIC1 peptides induced by a nitrate excess^39^. Importantly, the systemic regulation intrinsic to AoN remained unaffected by nitrate availability, as did the systemic control of CR development mediated by LalbCCR1 with respect to phosphate availability, once again suggesting that the AoDev pathway acts at a strict developmental level.

Both nodules and CRs serve as substantial carbon-sinks, primarily due to carbon demand of rhizobia within nodules and the carbon losses resulting from organic acid secretion by CRs. This developmental control is intricately regulated by shoot, in order to maintain optimal nodule or CR numbers. Common regulation by the AoDev pathway suggests a mutualization of carbon sink control and a potential recycling of genetic pathways. This raises several questions such as whether CR systemic regulation was duplicated from AoN in white lupin, and whether non-legume species harbor a systemic regulation of LR development independent of nutrient availability.

Indeed, the scenario is rendered more complex by recent findings in Medicago, elucidating the involvement of an MtCLE35 peptide perceived by the SUNN shoot LRR-RLK in the nitrate-mediated inhibition of nodulation, while MtCLE12/13-SUNN LRR-RLK orchestrates nitrate-independent systemic regulation of nodulation^40^. The interplay between N and P in governing both nodule and cluster root development is complex^18,38^, with numerous local and systemic control pathways, involving different peptide-LRR-RLK modules. However, the systemic AoDev pathway appears to be the dominant regulatory force^39^. Further studies will be needed to fully understand how systemic control of root system plasticity by the AoDev pathway is orchestrated and interacts with local components in legume and also in non-legume species.

## METHODS

### Plant material and cultivation

White lupin (*Lupinus albus*) cv. AMIGA (Florimond Desprez, France) and narrow-leaved lupin (*L. angustifolius*) cv. TANJIL (CSIRO, Australia) were used in this study. Seeds were germinated on a vermiculite substrate for 4 days, after which they were cultivated in 200 L hydroponic tanks containing the following well-aerated nutritive solution: 400 µM Ca(NO_3_), 54 µM MgSO_4_, 0.24 µM MnSO_4_, 0.1 µM ZnSO_4_, 0.018 µM CuSO_4_, 2.4 µM H_3_BO_3_, 0.03 µM Na_2_MoO_4_, 10 µM Fe-EDTA and either 200 µM K_2_SO_4_ for phosphate-deficient (-P) or 400 µM of KH_2_PO_4_ for phosphate-rich (+P) experiments. Growth chambers are set to a photoperiod of 16h light / 8h dark, 25,C day / 20,C night, 65% relative humidity and photon flux density of 200 µmol m^−2^ s^−1^. Grafting experiments were conducted at 7 days post-germination. The scion was prepared by trimming to a V-shape and inserted into a vertical slit created in the rootstock. Post-grafting, roots recovery rates ranged from 20 to 60% over the subsequent 3 to 7 days, contingent to the specific scion-rootstock pairing. Phenotyping analyses were conducted on 20-day-old plants. The number of LRs exhibiting clusters of rootlets, referred to as cluster roots (CR) in this study, were counted. Measurements of primary and LR lengths, as well as root dry weight, were also measured. Data were plotted and statistically analyzed using GraphPad Prism software 10.2. In order to visualize physiological activities of *ccr1-1* root systems, roots were spread on 0.8% agar plates containing either, 0.005% (m/v) bromocresol purple buffered in Tris-HCl pH 6 for testing proton excretion, or 0.013% (m/v) 5-Bromo-4-chloro-3-indolyl phosphate buffered in sodium acetate pH 5 for testing phosphatase activity, or 330µM bathophenanthroline disulfonic acide disodium salt, 100 µM FeNaEDTA for testing ferric reductase activity.

### EMS population and genetic screen

A large-scale mutagenesis was conducted on AMIGA seeds using 0.4% ethyl methanesulfonate (EMS) for 3 hours and deactivation with sodium thiosulfate 2.5% for 5 minutes. M1 seedlings were subsequently cultivated in the Cerience experimental fields (Poitiers, France) and the pods from each individual M1 plant were harvested, resulting in an EMS-mutagenized population of 5000 M2 batches. Finally, 36 seeds from each of 800 M2 batches were screened in P-rich medium to identify plants exhibiting constitutive cluster roots. Plants with the same constitutive cluster root phenotype were found in 4 independent batches and amplified. They were crossed for allelic test and back-crossed with AMIGA for the mapping by sequencing strategy. Pools of 50 to 90 F2 plants with a mutant phenotype (homozygous) and similarly sized pools of plants with a wild-type phenotype (WT batch) were harvested for DNA extraction and sequencing by Illumina HiSeq at Get-PlaGe core facility (INRAe, Toulouse, France). A mean coverage of 50X to 100X was obtained across samples. Cutadapt v1.15^41^ has been used to remove IlluminaTruseq adapter from the sequencing data and to remove bases with a quality score lower than 20, in both 5′ and 3′ end of the reads. Pairs of reads containing one read with a length lower than 35 have been discarded. We used BWA-MEM v0.7.17^42^ to map reads to the white lupin reference genome. Picard MarkDuplicates v2.20.1 (https://github.com/broadinstitute/picard) has been used to detect and remove PCR and Optical duplicates. We then used GATK HaplotypeCaller v4.1.4.1^43^ tool to call variants and snpEff 4.3t^44^ to annotate them. The duplicate free mapped reads have been used as input for the mutmap pipeline v2.1.2^45^.

### Microscopy

Root segments transitioning to CR were fixed with 4% paraformaldehyde for 120 min at room temperature under vacuum treatment and then washed twice for 2 min in 1X PBS before being embedded in 3% (w/v) agarose resin in PBS. Longitudinal root sections of 100 µm were cut with a vibrating microtome (Microcut H1200, BioRad). The sections were stained with 2 µg/mL 4,6 diamidino-2-phenylindole (DAPI). Fluorescence was observed using a ZEISS Axio Observer microscope, with a plan-apochromatic 20X/0.8 objective and the following filters: BP 325-390 nm for excitation and BP 445/50 for emission. Mosaic pictures were taken using the Apotome module. Images were captured with OrcaFlash (Hamamastu) controlled with the ZEISS Zen blue Software. In nodulation experiments, nodules were observed in dark field with the OLYMPUS SZX16 stereo microscope and images were taken with a DP72 camera.

### Structural and Phylogenetic analysis

AlphaFold structure prediction was performed using ColabFold v1.5.5: AlphaFold2 with MMseqs2^46,47^, providing the amino acid sequence of LalbCCR1 (LalbChr03g0025491). PyMol v2.5.4 was used to visualize and modify the protein structure to make the transmembrane domain apparent, and to indicate the positions of amino acid substitutions within the structure. Closest LalbCCR1 homologs from white lupin, narrow-leaved lupin, *Glycine max*, *Lotus japonicus*, *Medicago truncatula* and *Arabidopsis thaliana* were retrieved using NCBI blastp tool. Alignment of the kinase domain portion was conducted using MUSCLE^48^ alignment software on NGPhylogeny website with default settings, and the output was refined with JalView 2.11.3.3^49^ Phylogenetic analysis was performed using the PhyML/OneClick workflow with default settings on NGPhylogeny^50^. The resultant phylogenetic tree was generated using iTOl v6^51^. Synteny analysis utilized genome assemblies from *M. truncatula* A17 r5.0^52^, *G. max* Williams 82 v4.0^53^, *L. japonicus* MG20 v3.0^54^ and *L. albus* v1.0^30^. It was carried out using Easyfig 2.2.5^55^ with blastn and a minimum identity value for the blast at 0.7. The output of Easyfig was subsequently edited with Inkscape 1.2.1.

### Nodulation assays

For nodulation experiments plants were grown in Magenta GA-7 pot filled with leached and sterilized zeolite substrate (Siliz 14, Somez, France) supplied with a nutrient solution corresponding to the previously described P-rich medium but without nitrogen. Seeds were sterilized with calcium hypochlorite, germinated in Petri dishes and then transferred into Magenta pots. The *Bradyrhizobium lupini* MIAE428 strain (previously named LL13)^56^ was used. Inoculum was produced by cultivating the strain in modified yeast mannitol (YM) medium (mannitol 10 g/L, yeast extract 1 g/L, K_2_HPO_4_ 0.5 g/L, NaCl 50 mg/L magnesium sulfate 7H_2_O 100 mg/L, calcium chloride 40 mg/L, glutamic acid 0.43 g/L, FeCl_3_ 4 mg/L) supplemented with nalidixic acid 20 µg/L, in the dark for 4 days at 28 °C. One mL inoculum was applied one week after the seedlings were transferred to the pots or one week after grafting. Nodule numbers per plant were assessed and the leaf chlorophyll content was indirectly estimated using a Chlorophyll meter SPAD (Konica-Minolta) on the third youngest leaf. Data were plotted and statistically analyzed using GraphPad Prism software 10.2.

### Gene expression analysis

Developmental temporal transcriptome. The sampling began (T0) on eight-day-old plants grown in P-deficient conditions. A total of eight 1 cm-long transitioning CR segments from four independently grown plants was sampled, at a distance of 1 cm from the primary root, in the upper part of the root system where LRs are transitioning to CR, every 12 h for 5 days, covering the entire rootlet developmental process (T0 to T132). As a control, 1 cm-long lateral root segments not transitioning to CR were collected. Four biological replications were produced for each experiment. Total RNA was extracted from all frozen samples using the Direct-zol RNA MiniPrep kit (Zymo Research) according to the manufacturer’s recommendations. A total of 52 independent root RNA-seq libraries were constructed and sequenced at Get-PlaGe core facility (INRAe, Toulouse, France). The Illumina TruSeq Stranded mRNA Sample Preparation Kit (Illumina Inc.) was used according to the manufacturer’s protocol. Paired-end sequencing was performed, generating 2 x 150 bp reads using TruSeq SBS kit v3 sequencing chemistry on an Illumina NovaSeq instrument. Raw reads were cleaned using Cutadapt v1.15^41^, by removing bases with a quality score lower than 30, in both 5′ and 3′ end of the reads, as well as TruSeq Illumina adapters. Pairs of reads containing one read with a length lower than 35 have been discarded. The quality-checked RNA-seq reads were mapped on the white lupin genome reference using Hisat2 v2.1.0, with the following parameters “--rna-strandness RF --dta”. Transcripts were assembled and quantified using Stringtie v1.3.4d with the options “--rf -e -B -u - M 1”.

*ccr1-1* mutant transcriptome. All LRs of four ten-day-old plants grown in P-deficient conditions were harvested from the upper part of the root system, corresponding to the zone where LRs are transitioning to CR. Eight independent RNA-seq libraries were constructed and processed as described for developmental temporal transcriptome. Normalization, differential expression and gene ontology enrichment analysis were performed using the DIANEbeta R package^57^ (https://shinyapps.southgreen.fr/app/dianebeta) v1.1.0.1. The TCC R package with the “tmm” normalization method was used, with prior removal of differentially expressed genes. For each analysis, Log2(fold change) and False Discovery Rate adjusted p-value (FDR) were provided in the text and figure legends. SRPlot online^58^ was used for generating the PCA, heatmap, and the GO plots, and GraphPad Prism software 10.2 for the statistical analysis and kinetic expression data plotting. The GO terms used for enrichment are available for download at https://www.whitelupin.fr/download.html.

### RT-qPCR experiments

For *LalbCCR1* expression in different organs, 11-day-old plants grown under either P-deficient or P-rich conditions were sampled. Samples included for lateral roots (LR), root apical meristem (RAM), shoot apical meristem (SAM), leaf, petiole and hypocotyl. Cluster root (CR) samples were collected exclusively from plants grown on P-deficient conditions. CRs were collected from the upper part of the root system, while LRs were collected below. The apices of LR and CR were removed. Petioles and leaves were collected from the second leaf. Each sample contained tissues from 3 individual plants and 3 biological replicates were collected for each plant part. For experiments confirming transcriptomic data in another allele besides *ccr1-1*, samples were collected from the root system of *ccr1-2* mutant plants following the same protocol as for the transcriptomic study. For the grafted plants, this protocol was applied to roots recovering after the grafting operation. At least five biological replicates were collected and analyzed in each experiment. Total RNA was extracted using the Direct-zol RNA MiniPrep kit (Zymo Research) according to the manufacturer’s recommendations. RNA concentration was measured on a NanoDrop (ND1000) spectrophotometer. Poly(dT) cDNAs were synthetized from 2 µg total RNA using the RevertAid First Strand cDNA Synthesis (ThermoFisher). Gene expression was measured by quantitative Real Time-Polymerase Chain Reaction (qRT-PCR) (LightCycler 480, Roche Diagnostics) using the SYBR Premix Ex Taq (Tli RNaseH, Takara, Clontech). Expression levels were normalized to a putative initiation factor *LalbEIF-4* (*Lalb_Chr07g0195211*) or to a *LalbPolyubiquitin* (*Lalb_Chr06g0164891*). Two technical replicates were performed for all qRT-PCR experiments. Specific primer pairs are described on the Supplementary Table 10. Relative gene expression levels were calculated according to the ΔΔCt method, using LR (for organ expression), or WT and WT/WT, samples for *ccr1-2* and grafted plants respectively.

### Data availability

FASTQ raw sequence files are available at NCBI under the Bioproject number PRJNA1124865 for the temporal RNAseq (Sequence Read Archive accession numbers SAMN41865670-82) and number PRJNA1125199 for the *ccr1-1* RNAseq (Sequence Read Archive accession numbers SRR29446565-79).

*Reviewer links for temporal dataset:* https://dataview.ncbi.nlm.nih.gov/object/PRJNA1124865?reviewer=851s8ptss62fl7h178k2j5f8bu *and for ccr1-1 dataset:* https://dataview.ncbi.nlm.nih.gov/object/PRJNA1125199?reviewer=m1lhh43j385of3s7hgrnsd7ucp

## Acknowledgements

We thank Nathalie Harzic (Cerience, Poitiers) for the white lupin EMS mutagenesis and agricultural support. We thank Carine Alcon from the PHIV facility (Plateforme d’Histocytologie et d’Imagerie cellulaire Végétale of IPSiM lab), for her valuable comments and technical assistance with microscopy imaging. We thank Cécile Revellin (UMR Agroécologie, INRAE Dijon) for the gift of MIAE428 *Bradyrhizobium lupinus* strain. We also wish to thank Valérie Hocher and Darius T. Nzepang (IRD, Montpellier) for technical help in setting up nodulation protocols for white lupin. We thank Lars Kamphuis and Karam Singh (CSIRO, Australia) for providing narrow-leafed lupin TANJIL seeds.

## Author contributions

FD and FG performed the initial genetic screen and subsequent genetic analysis. FD, AB, LM, VF, CMC, IV performed phenotypic analysis. LM, CMC, FD and AB performed grafting experiments and expression analysis. BH generated the temporal RNAseq dataset and FD the *ccr1* dataset. EIA performed the *CCR1* expression analysis. AS, HP and LM performed bioinformatic analyses. LM analyzed the data. LM and FD generated the figures. LM and BP conceived the project, obtained funding and wrote the article.

