## Extended data figures for "Autoregulation of cluster root and nodule development by white lupin CCR1 receptor-like kinase"

**Extended data Figure 1. Phenotypic data of *L. albus ccr1* mutants**

a) Representative images of the four allelic *ccr1* mutants at 20-days-old, grown on either phosphate-rich medium (+P) or phosphate-free medium (-P). Scale bar = 1 cm.

b) Length of the primary root of the four allelic *ccr1* mutants at 20-days-old, grown on either phosphate-rich medium (+P) or phosphate -free medium (-P). Statistical analysis was performed using two-way ANOVA with Tuckey correction, n = 12, $p < 0.05$.

c) Physiological activity of the root system of the *ccr1-1* mutant. Root systems were spread on agar plates containing bromocresol purple for testing proton excretion, or 5-Bromo-4-chloro-3-indolyl phosphate for testing phosphatase activity, or bathophenanthroline di-sulfonic acid disodium salt for testing ferric reductase activity. Scale bar = 1 cm.

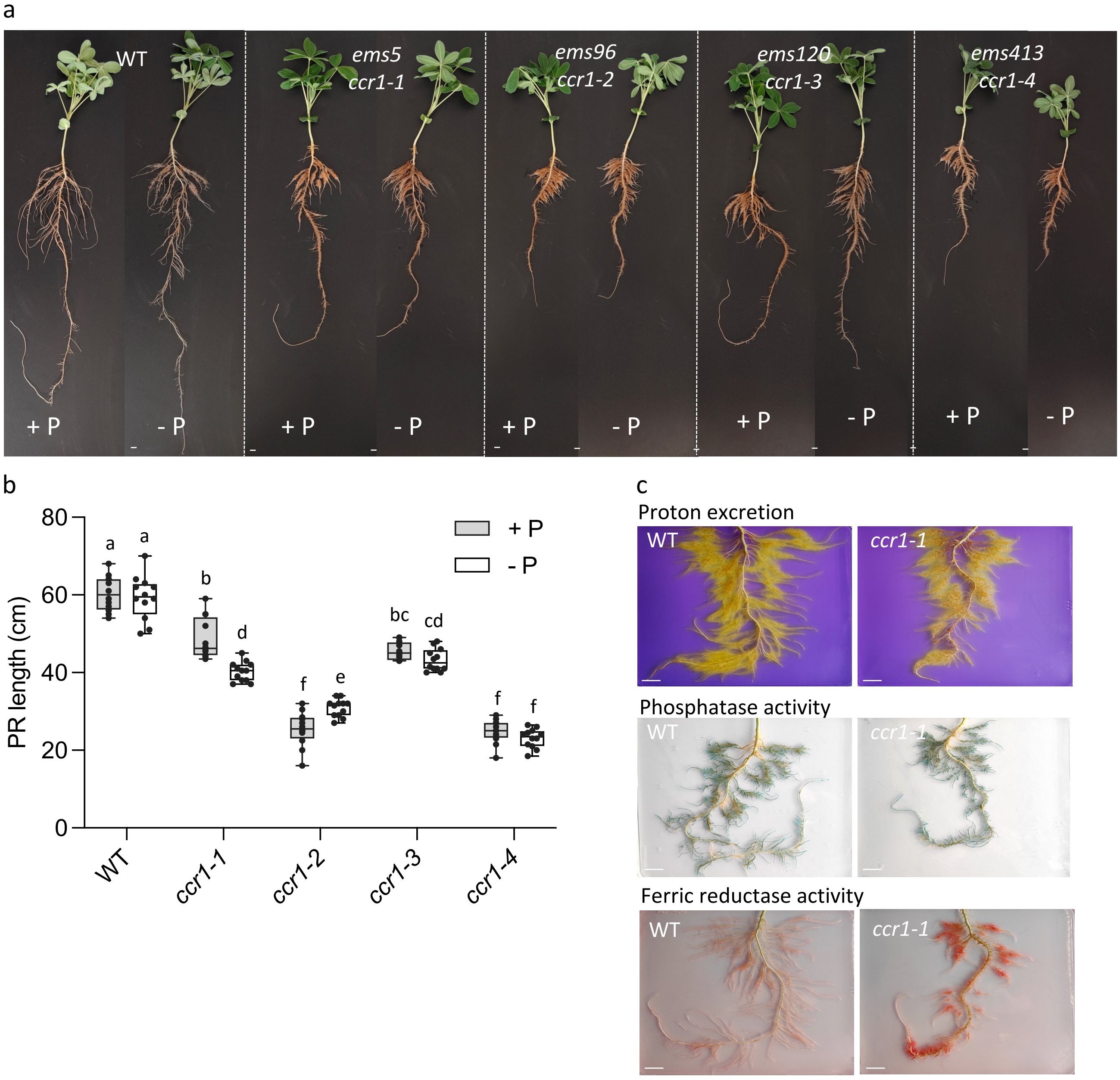

**Extended data Figure 2. Genetic and molecular data for *L. albus ccr1* mutants alleles**

a) Molecular data of the causal SNPs in the four *ccr1* alleles. F2 genetic segregation for the WT and *ccr1* phenotypes showed a 3:1 ration consistent with recessive alleles. Results of allelism tests indicated a unique complementation group.

b) Phenotyping and sequencing of WT and *ccr1-1* plants from a second backcross (*ccr1-1* x WT) identifying the causal SNP G2099>A at heterozygotous or homozygotous level in the descendants.

c) Mapping by BSA-seq analysis of SNPs associated with mutant or wild-type phenotypes from backcrosses with the four *ccr1* alleles. Significant *ccr1*-mutant-associated-SNPs were observed at the beginning of chromosom 3 for the four alleles in the mutant-phenotype bulk.

a

| Gene | Alleles | | Mutation | | F2 segregation | | Complementation group |
|---|---|---|---|---|---|---|---|
| | ems name | mutant name | nucleotide level | amino acid level | WT : *ccr1* | P>0.05 * | |
| Lalb_Chr03g0025491 | ems5 | *ccr1-1* | G2099>A | G700>E | 254 : 84 | 3 : 1 * | a |
| | ems96 | *ccr1-2* | G1370>A | W457>stop | 295 : 96 | 3 : 1 * | |
| | ems120 | *ccr1-3* | C2437>T | H813>Y | 311 : 94 | 3 : 1 * | |
| | ems413 | *ccr1-4* | C2437>T | H813>Y | 291 : 81 | 3 : 1 * | |

b

| BC2 *ccr1-1* x WT | Phenotype | | Causal SNP | | Genotype | P>0.05 * |
|---|---|---|---|---|---|---|
| 42 | WT | 32 | (G /G) | 10 | (+/+) | 1/4 |
| | | | (G /A) | 22 | (+/-) | 1/2 |
| | *ccr1* | 10 | (A /A) | 10 | (-/-) | 1/4 |

c

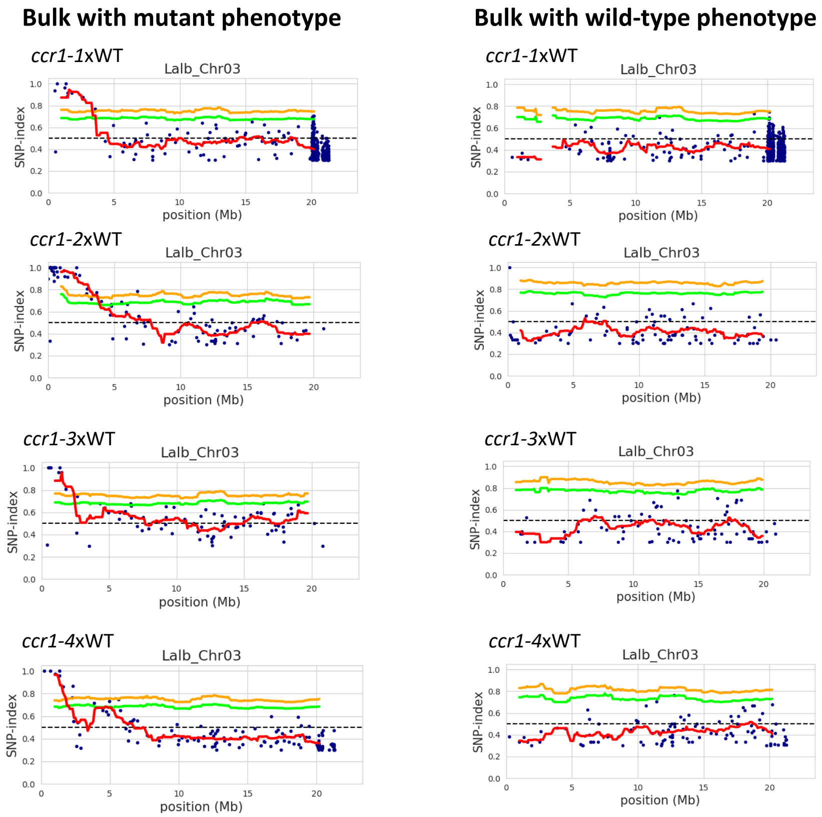

**Extended data Figure 3. Hypernodulation phenotypes of *L. albus ccr1-1 and ccr1-2* mutants.**

a) Representative images of the aerial parts of non-inoculated wild-type and *ccr1-1* mutant plants, grown on P-rich, N-deprivated medium. Scale bar = 1 cm.

b) Representative images of the roots of non-inoculated wild-type and *ccr1-1* mutant plants grown on P-rich, N-deprivated medium. Scale bar = 0.5 cm.

c) Representative images of the aerial parts of wild-type and *ccr1-1* mutant plants grown on P-rich, N-deprivated medium and inoculated with *Bradyrhizobia lupini*. Scale bar = 1 cm.

d) Representative images of the roots of wild-type and *ccr1-1* mutant plants grown on P-rich, N-deprivated medium and inoculated with *Bradyrhizobia lupini*. Scale bar = 0.5 cm.

e) SPAD measurements of leaf color in wild-type and *ccr1-1* mutant plants, either non- inoculated or inoculated with *Bradyrhizobia lupin.* The SPAD values of the inoculated plants are higher than those of the non-inoculated ones, indicating efficient nodulation for nitrogen nutrition. One-way ANOVA test, n = 40 to 48, $p < 0.0001$.

f) Representative image of cut nodules showing their reddish color due to the presence of leghemoglobin. Scale bar = 0.2 cm.

g) Representative image of wild-type and *ccr1-2* mutant plants grown in Magenta boxes on P-rich, N-deprivated medium and inoculated with *Bradyrhizobium lupini*. Scale bar = 1 cm.

h) Representative image of wild-type and *ccr1-2* mutant CRs from one plant grown in Magenta boxes on P-rich, N-deprivated medium and inoculated with *Bradyrhizobium lupini*.

i) Representative image of cluster of nodules from wild-type and *ccr1-2* mutant plants grown in Magenta boxes on P-rich, N-deprivated medium and inoculated with *Bradyrhizobium lupini*. Scale bar = 0.2 cm.

j) Nodule number per plant for wild-type, *ccr1-1* and *ccr1-2* mutant plants grown in Magenta boxes on P-rich, N-deprivated medium and inoculated with *Bradyrhizobium lupini*. One-way ANOVA test, n=10, ns = non statistically different, ***p-value < 0.001.

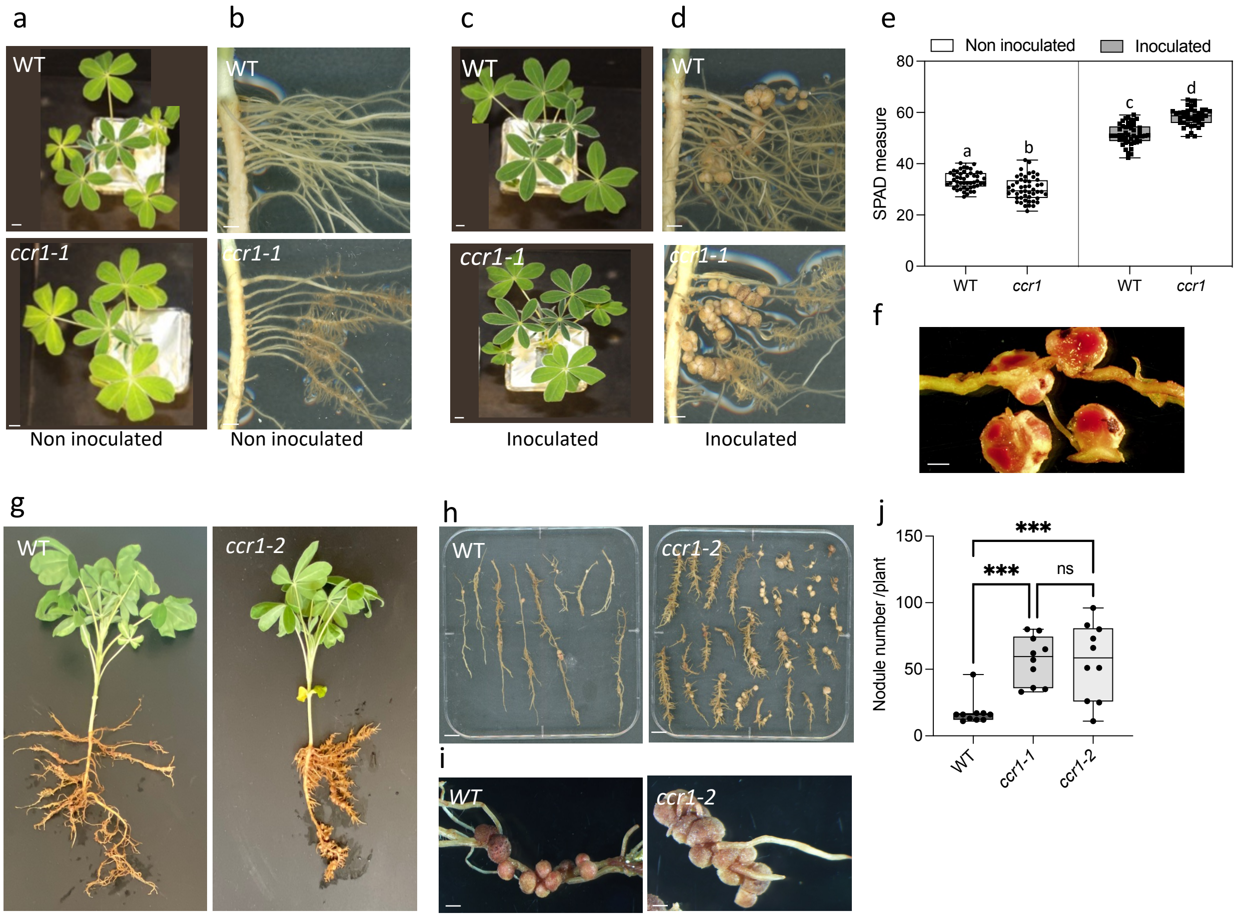

**Extended data Figure 4. Expression analysis of *LalbCCR1* in different organs of white lupin plants.**

Shoot apical meristem (SAM), leaf, petiole, hypocotyl, root apical meristem (RAM), lateral root (LR; secondary roots without rootlets) were sampled from plants grown on either phosphate-rich medium (+P) or phosphate-free medium (-P). Cluster roots (CR; secondary roots with cluster of rootlets) were sampled from plants grown on phosphate-free medium (-P) only since there was no CR on plants grown on phosphate-rich medium (+P) (nd: no data). Each sample contained tissues from three individual plants, and three biological replicates were collected for each plant part. Mean with standard errors are shown on the graph.

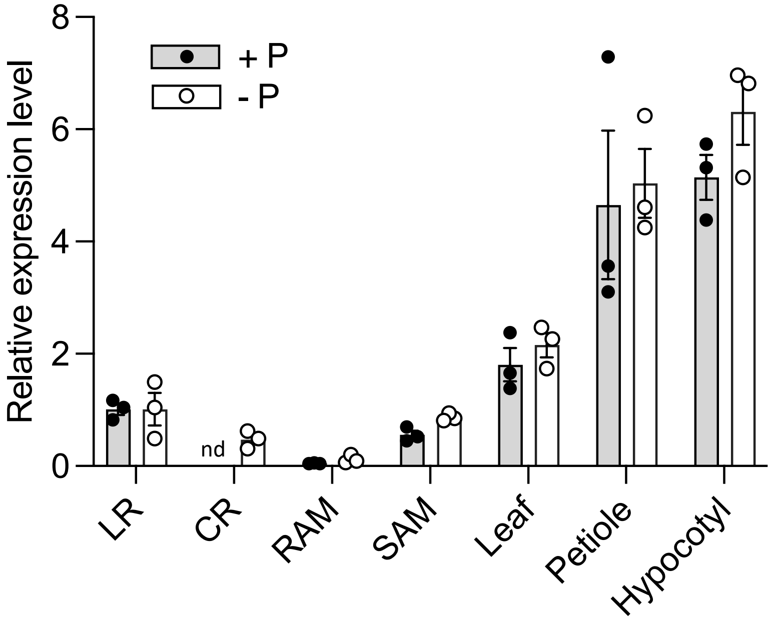

**Extended data Figure 5. Grafting experiments with the allele *Lalbccr1-2.***

a) Representative images illustrating rootstock phenotypes of grafted plants using wild-tpe (WT) plants as rootstocks and WT or *ccr1-2* mutant as scions. The *ccr1-2* scion triggers the formation of numerous CRs on WT rootstock. Legends of the images indicate the scion/rootstock combination. Scale bar = 1 cm.

b) Representative images illustrating rootstock phenotypes of grafted plants using *ccr1-2* mutant plants as rootstocks and WT or *ccr1-2* mutant as scions. The WT scion inhibits the formation of CRs on *ccr1-2* rootstock. Legends of the images indicate the scion/rootstock combination. Scale bar = 1 cm.

c) Representative images illustrating rootstock phenotypes of grafted plants using NLL (*L. angustifolius*) as rootstocks and NLL or the mutant *ccr1-2* as scions. The *ccr1-2* scion triggers the emergence of numerous tertiary roots on NLL rootstock. Legends of the images indicate the scion/rootstock combination. Scale bar = 1 cm.

d) Representative images illustrating rootstock phenotypes of grafted plants using *ccr1-2* as rootstocks and NLL or the mutant *ccr1-2* as scions. The NLL scion significantly inhibits the formation of CRs on rootstock. Legends of the images indicate the scion/rootstock combination. Scale bar = 1 cm.

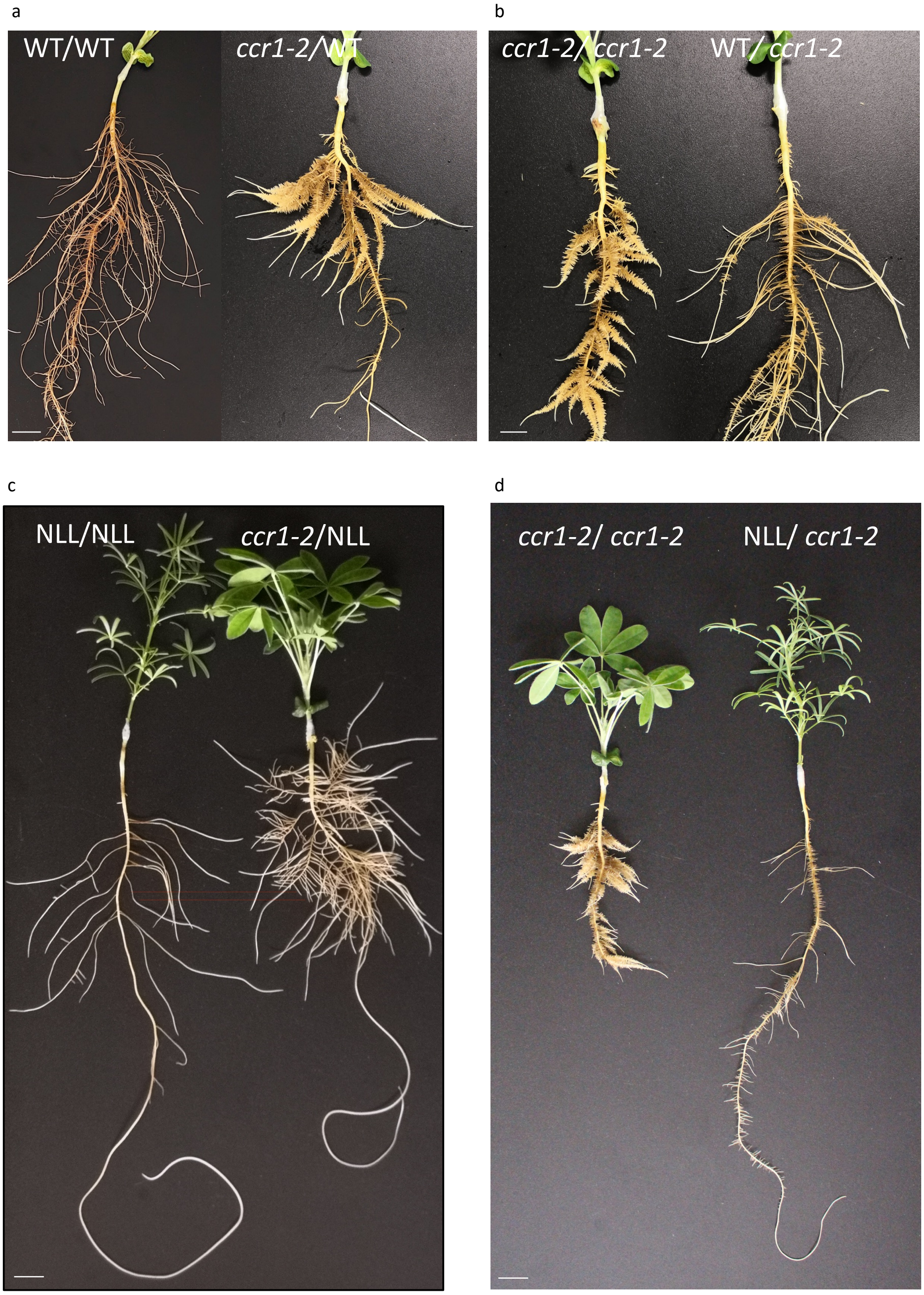

**Extended data Figure 6. GO enrichment analysis of down-regulated genes from the temporal developmental  RNAseq dataset.**

Down-regulated genes between LR and timepoints T024-036-048 (Absolute Log2(fold change) > 2, FDR < 0.05 ) and presenting  counts > 10 are displayed. The GO terms retrieved are primarly associated with stress responses and cell-wall modifications. The analysis was performed  using DIANEbeta R package.

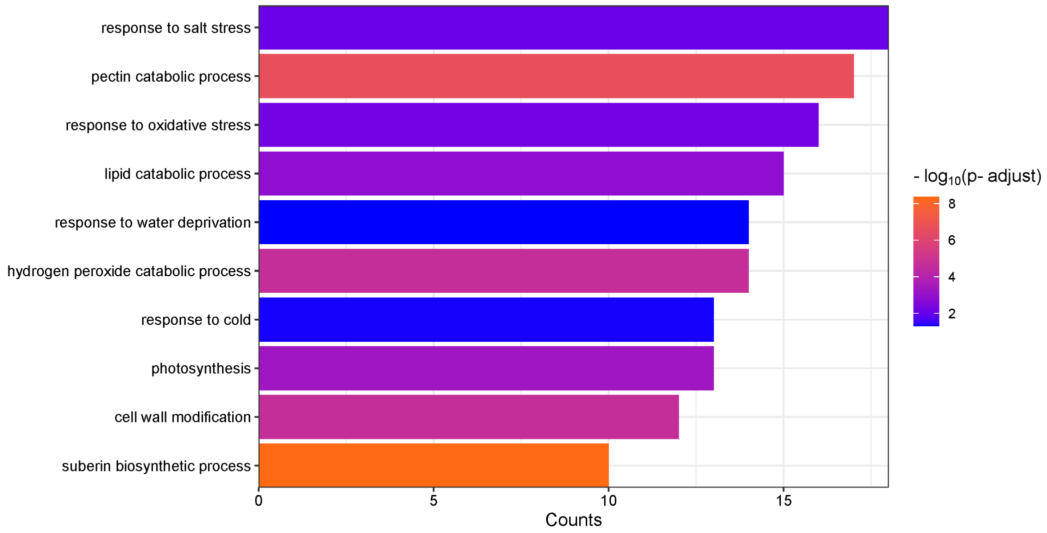

**Extended data Figure 7. Analysis of differentially regulated genes between *ccr1-1* and wild-type roots.**

a-b) GO term analysis presenting counts > 5. The DEG analysis was performed using DIANEbeta R package. Absolute Log2(fold change) > 1, FDR < 0.05. a) up-regulated genes; b) down-regulated genes. Underlying data can be found in Supplementary Table 8.

c-d) RT-qPCR analysis of the expression levels of selected genes in roots of *ccr1-1* and *ccr1-2* mutants compared to wild-type. The roots were sampled at the same developmental stage than for the RNAseq *ccr1-1*/WT samples. Error bars represent mean ± SE, and statistical analysis was performed using Two-tailed Mann Whitney tests: *p-value <0.05; **p-value <0.01; ***p-value <0.001; ****p-value<0.0001.

a)

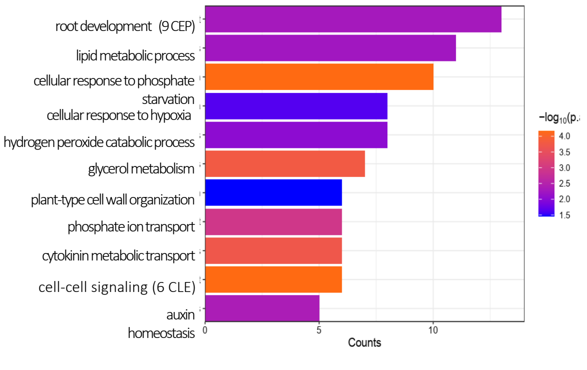


b)

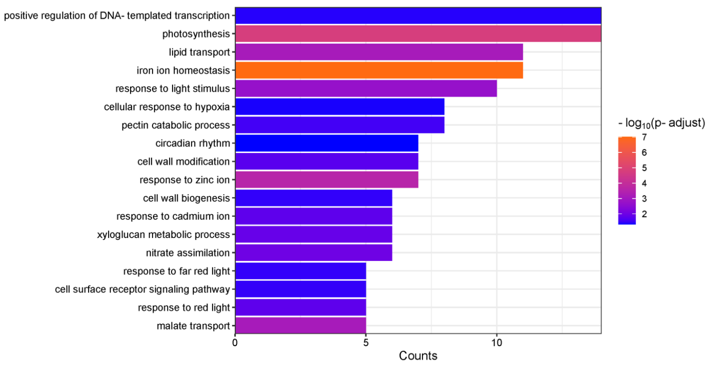


c)

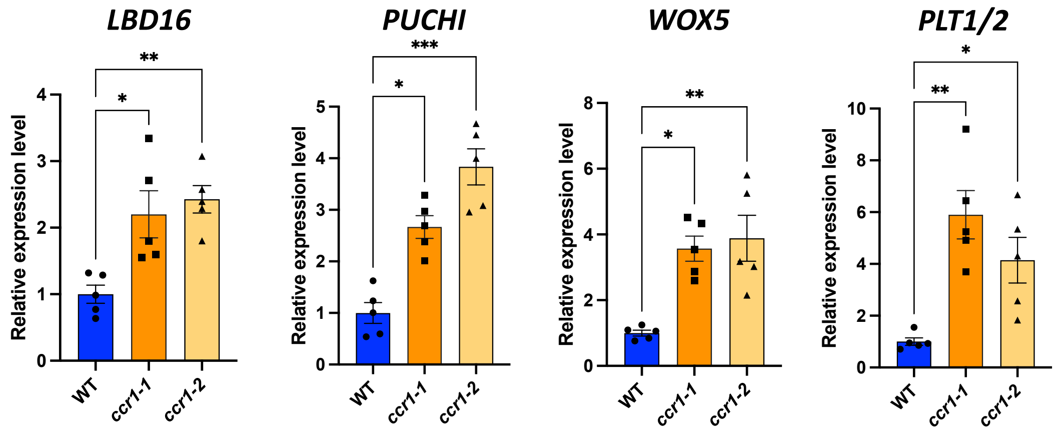


d)

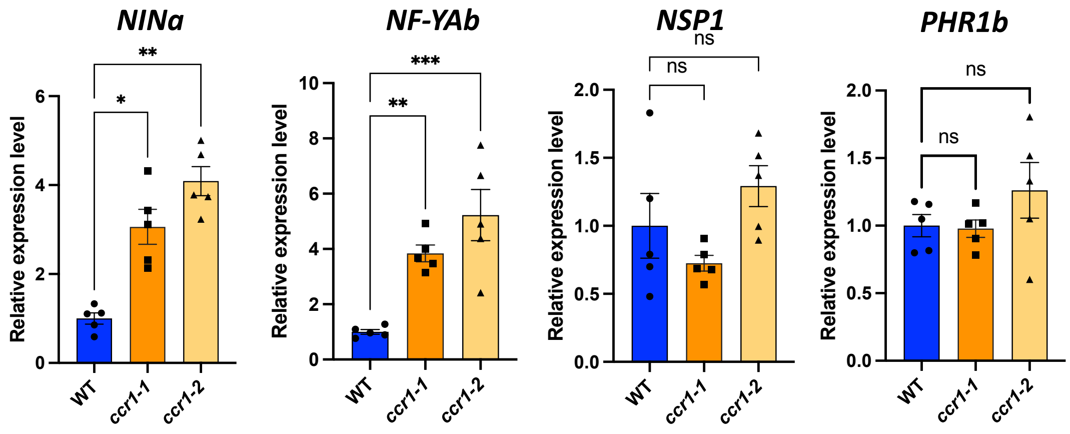
